# The visual social environment affects non-additively neighbor spacing and interaction time in zebrafish

**DOI:** 10.1101/511972

**Authors:** Diana Pita, Esteban Fernández-Juricic

## Abstract

Many fish form schools and visually track the position of their neighbors in a 3D environment. In this study, we assessed whether zebrafish modified their spacing behavior and interaction time in an additive or multiplicative way relative to multiple sources of visual social information using video playbacks. We simultaneously manipulated: (a) the magnitude of the social cues (by varying the size of the virtual fish), (b) the level of social risk (low, high based on the position of the virtual fish in the water column), and (c) the perceived depth of the social cues (visual horizon absent or present). Each of these factors independently affected spacing behavior (zebrafish increased the separation distance with larger virtual fish, under lower visual social risk, and when depth perception was enhanced), but they did not affect interaction time. However, some of these factors interacted significantly, such that their effects on social behavior depended on each other. For instance, zebrafish decreased their separation distance under high vs. low risk conditions when the virtual fish was the same or smaller size, but this risk effect disappeared with larger virtual fish likely to avoid aggression. Also, zebrafish increased their separation distance when perceived depth was enhanced under low risk, but the perceived depth effect became less pronounced under high risk probably due to dilution effects. Overall, the effects of certain visual social parameters depend on the intensity of other visual social parameters, ultimately tuning up or down different social behavioral responses. We discuss the implications for the spatial organization of fish schools.

**Significance Statement:** Many fish form schools and visually track the position of their neighbors in a 3D environment. We found that zebrafish consider multiple visual social sources of information simultaneously to modify their neighbor distance. Thus, their spacing behavior appears to follow multiplicative rules, whereby the spacing response to a visual social parameter depend on the intensity of a different visual social parameter.

## Introduction

In animal groups, an individual’s interaction range represents the area over which it can acquire and respond to social information (Sumpter 2010). The interaction range can be characterized as a circular zone surrounding the individual (Krause and Ruxton 2002). In the outer area (i.e., attraction zone), individuals move closer to group mates; in the middle area (i.e., alignment zone), individuals orient in the same direction as group mates; and in the center area (i.e., repulsion zone), individuals move away from group mates (Krause and Ruxton 2002). Ultimately, the size of the interaction range defines the number of group mates that an individual can gather social information from (Fernandez-Juricic et al. 2007).

An important factor that can limit the distance over which social information is transferred is the perceptual ability of the receiver (Strandburg-Peshkin et al. 2013; Pita et al. 2016). In the case of visual information, the limits of spatial resolving power or visual acuity (i.e. ability to resolve two objects as separate entities) can constrain the size of the interaction range (Pita et al., 2016). Changes in the size of the interaction range are expected to change an individual’s behavior to maintain its ability to resolve social information (Kimbell and Morrell 2015). For instance, when social information becomes more difficult to gather, individuals can shorten their neighbor distances (Dawkins and Woodington 1997), lengthen the duration of their interactions with group mates (Fernández-Juricic et al. 2004; Fernandez-Juricic et al. 2007), or use regions of the visual field with high visual acuity (i.e., centers of acute vision; Butler and Fernández-Juricic 2014) to gather social information.

Animals use social cues to regulate their social interactions (Pritchard et al. 2001, Fernández-Juricic and Kacelnik 2004; Abaid et al. 2012). The availability of virtual playbacks and robotic animals has allowed for manipulating these social cues to assess their role in tuning social behavior (e.g., Gerlai 2017, Bierbach et al. 2018; Kim et al. 2018). Several studies have manipulated single social cue types at a time (e.g., Butail et al. 2015, Bartolini et al. 2016); however, we know relatively less about the effects of *simultaneous* different social cues types on an individual’s ability to gather social information and respond to variations in its visual social environment (but see Larsch & Baier 2018; Lemasson et al. 2018). This is particularly relevant because understanding how animals respond to variations in the social environment can allow us to identify the rules underlying social interactions (Arganda et al. 2012). For instance, two different visual social cues (e.g., color and size) can act in an additive (i.e., the combined effect of color and size is equal to the sum of each effect taken individually) or multiplicative (i.e., the effect of color cues depends on the size cues) way. Furthermore, we can explore how these rules may be affected by environmental change, leading potentially to the disruption of social interactions and changes in the spatial arrangement of groups. For example, increased water turbidity can reduce the transfer of visual social information between fish, resulting in fewer social interactions and smaller group sizes (Borner et al. 2015).

In this study, we assessed how individuals modified their social spacing behavior and social interaction time relative to three types of simultaneous changes in the visual social environment: (a) the magnitude of the social cues (by manipulating the size of the conspecific eyes and body stripes), (b) the level of visual social risk (by manipulating the position of the conspecific in the water column), and (c) the perceived depth of the social cues (by manipulating the absence/presence of the visual horizon). We used zebrafish as our model species because they are highly social animals that depend on visual information during social interactions (Miklosi and Andrew 2006; Saverino and Gerlai 2008). We manipulated the visual social environment using a video playback technique following Balzarini et al. (2017) and Ingley et al. (2015). We considered this playback technique to be an appropriate method because zebrafish are responsive to video playbacks (Saverino and Gerlai 2008; Stowers et al. 2017), reacting to animated conspecifics in the same way they do to live conspecifics (Qin et al. 2014; Gerlai 2017). We only considered interactions with a single conspecific as a first attempt to identify some of the simplest rules underlying pairwise social interactions. Future studies should consider how these rules scale up in larger groups (e.g., Lemasson et al. 2018).

### Hypotheses and predictions

We first present predictions for single effects and then for their interactions.

An individual’s ability to resolve social cues from a neighbor (e.g., variations in coloration, body patterns, etc.) decreases with distance (Fernández-Juricic and Kowalski 2011). To simulate changes in the magnitude of the social cues over distance, we manipulated the size of the conspecific social cues (i.e., eye diameter and width of body stripes), which scaled directly with conspecific body size. We predicted that as the magnitude of the social cues decreased, individuals would move closer to the conspecific to enhance their ability to resolve them (Pita et al. 2015) and/or increase the amount of time spent interacting with the conspecific (Fernández-Juricic et al. 2004; Fernandez-Juricic et al. 2007).

In zebrafish, the vertical position of the fish in the water column varies with the level of risk (Bishop et al. 2016; Thompson et al. 2016; Oliveira et al. 2017)). Zebrafish located at the bottom of the water column (i.e., low vertical spatial position) are associated with higher risk environments (e.g., predator presence; Kalueff et al., 2013; Oliveira et al., 2017). We therefore predicted that when the conspecific was at the bottom of the water column, individuals would move closer to the conspecific and/or increase the duration of their interaction to obtain more information about the perceived threat (Pham et al. 2012).

When animals assess the size of an object, they consider the visual angle or angular size (i.e., amount of space that the object takes up on the retina; Douglas, 1996; Sperandio and Chouinard, 2015). However, distance can complicate this assessment, as objects that are far away have a reduced visual angle and consequently appear smaller than they really are. To compensate for this, animals have evolved various mechanisms to improve depth perception (Howard 2012; Nityananda and Read 2017), ultimately allowing them to establish size constancy (i.e. ability to perceive the relative size of an object independent of its angular size; Douglas et al., 1988; Frech et al., 2012; Zeil, 2000). One of the cues that facilitates depth perception is the presence of a visual horizon, which serves as a salient referential cue that can offer information about object orientation and distance (Layne et al. 1997; Caballero et al. 2015). In their natural habitats, zebrafish have a visual horizon defined by a variety of environmental substrates (Zimmermann et al. 2018). We manipulated the presence/absence of a virtual visual horizon in our video playbacks. We predicted that when the horizon was absent, individuals would move closer to the conspecific and/or increase the duration of the social interaction due to the lack of referential information and the need to better assess its relative size and location, compared to a situation in which the horizon was present.

Because we manipulated these three factors simultaneously, we were able to assess interaction effects. The conceptual frameworks we used allowed us to generate predictions for two-way interactions between pairs of the three factors considered (see also statistical analyses). First, we predicted a significant interaction between the magnitude of the social cues and the level of social risk, whereby individuals would increase their separation distance and reduce social interaction time particularly when exposed to a low magnitude social cue (i.e. small conspecifics) at the bottom of the water column. We hypothesized that individuals may experience higher risk when joining a group composed of smaller conspecifics in a high-risk scenario because their larger size would make them more conspicuous to potential predators (Krause and Ruxton 2002; Croft et al. 2006; Rodgers et al. 2015).

Second, we predicted a significant interaction effect between the magnitude and perceived depth of social cues. When the horizon was absent, zebrafish would likely perceive conspecifics according to their absolute size due to the lack of referential information, leading to conspecifics with high and low magnitude social cues appearing as smaller or larger, respectively, than conspecifics displayed with social cues of intermediate magnitude. In these scenarios (small/large social cues with horizon absent), we predicted zebrafish would increase their separation distance and reduce their interaction time to minimize the costs of interacting with differently sized group mates (Nakayama et al. 2009). For example, body size correlates with dominance in teleost fish and individuals of larger body size are more likely to win fighting contests (Arnott and Elwood 2009), while individuals of smaller body size are more likely to pay a cost for fighting (Hoare et al. 2000). On the other hand, when the horizon was present, individuals should perceive conspecifics according to their relative size (i.e., a small conspecific would be perceived to be far away compared to a large conspecific) because of an enhanced ability to perceived depth.

Third, we predicted an interaction between the level of visual social risk and the perceived depth, whereby individuals would move closer to the conspecific and increase their interaction time particularly when the conspecific is at the bottom of the water column and the horizon is present. We hypothesized that the closer the conspecific is to the horizon, the more pronounced the enhanced depth perception benefits would be, and consequently the perception of a high-risk scenario.

## Methods

We used 10 adult zebrafish (wildtype AB genetic strain) acquired from The Zebrafish International Resource Center (ZIRC, Eugene, Oregon, USA). Before the experiment, zebrafish were housed together in a 12-gallon glass stock tank (20.25’’ L x 10.5” W x 12.63” H) maintained with a recirculating water filter (Aqueon Power Filter ®) and exposed to a 16-hour light: 8-hour dark cycle. Water quality checks were preformed daily (pH, temperature) and weekly (ammonia, nitrates, nitrites, chlorine) to ensure appropriate housing conditions. We fed zebrafish daily with commercial fish flakes (Tetramin ^®^ Tropical Flakes).

Before the initiation of the experiment, zebrafish were transferred from their stock tank into individual 2.5-gallon tanks (12” L x 6” W x 8” H) with air stones to provide aeration. To reduce stress behaviors associated with exposure to a novel tank environment, each zebrafish was tested within its individual 2.5-gallon home tank. During treatment exposure, the home tank was placed directly against the screen of a computer monitor (Acer LCD monitor, model v176L) displaying the animated treatment video (supplementary material, Figure A1). The videos were generated using the open source software anyfish (http://swordtail.tamu.edu/anyfish/) which allows the user to create realistic 3D animated fish stimuli (Ingley et al. 2015). We used a photograph of an adult female wildtype zebrafish as a model for our animated stimulus. A female zebrafish was utilized because in other studies it has been shown that males are less likely to engage in interactions with other males, but females are preferred by both sexes (Ruhl and McRobert 2005). Following previous zebrafish video playback studies, we used one virtual exemplar per treatment (Saverino and Gerlai 2008; Gerlai et al. 2009; Qin et al. 2014). By modifying the x, y and z positions of the animated stimulus with user-defined key frames, we designed the virtual conspecific to swim back and forth, horizontally across the screen. The individual frames were then combined to generate a 60 frame per second video to simulate a conspecific swimming at approximately 5 cm/second which is the approximate speed maintained by zebrafish when schooling (Miller and Gerlai 2012). The flicker rate of the animation was designed to simulate smooth continuous movement in the eyes of the zebrafish as the critical flicker fusion frequency of many fish are thought to be within the range of 14-67 Hz (Lythgoe 1979).

We positioned top- and side-view cameras (JVC Everio GZ-MG330-HU camcorder) to record the behavior of the zebrafish throughout the trial (supplementary material, Figure A1). We synchronized both camera views with a portable DVR (Night Owl, H264-4 channel DVR) to allow for the determination of the 3D position of the zebrafish. The 6” sides and the 12” bottom of the tank were covered with opaque white tape and the experimental arena, consisting of a table that supported the tank and computer monitor were surrounded by opaque white curtains to reduce visual distractions during trials. Before each trial, there was a 10-min acclimation period followed by the 3-min treatment video that provided 1 min of continuous conspecific exposure. The first minute of the treatment video displayed only the virtual environment followed by 1 min of the conspecific swimming within in the virtual environment, while the last minute displayed again the virtual environment without the conspecific. Light levels at the surface of the water (mean ± SE; 1072 ± 1.35 lux) and temperature levels (mean ± SE; 74 ± 0.07 °F) remained constant throughout the experiment.

We exposed zebrafish (4.5 ± 0.15 cm body length; mean ± SE) to a total of 12 treatments, represented by the combination of 3 categorical independent variables: magnitude of the social cues (3 levels: low, intermediate, high), level of visual risk (2 levels: low, high) and perceived depth (2 levels: low, high). Each zebrafish was exposed to one treatment per day, allowing for approximately 24 hours between each successive trial, over the course of 12 days. Additionally, we randomized the testing order of the zebrafish each day, in addition to the order of each zebrafish’s series of treatment exposures.

Videos were coded manually and blind by an individual unaware of the experimental question or design. First, we used the program idTracker (http://www.idtracker.es/) to divide each video into frames (30 frames/s). Following this procedure, we used ImageJ (https://imagej.nih.gov/ij/) to measure the visual separation distance between the eye of the zebrafish and the body of the animated conspecific. We chose to manually code the videos (see also Delcourt and Poncin, 2012) because other fully-automated tracking programs tend to only measure the distance between the center of the zebrafish and the computer screen, but we intended our measurements to be based more sensory specific, as the information on distances to objects is gathered by the eye. An example of the movement trajectories we generated is in the supplementary material (Fig. A2). By using 3D measurements obtained from manual coding, we were able to calculate the position of the conspecific within the zebrafish’s visual field following (Pita et al. 2015) to determine if the conspecific was aligned with high or low acuity retinal regions. High acuity retinal regions are expected to provide finer scale spatial information compared to low acuity retinal regions (Pettigrew et al. 1988). In the case of zebrafish, a previous study found that each eye has one center of acute vision (enlargement of the retinal tissue due to the higher density of cone photoreceptors/reginal ganglion cells or *area*) that projects fronto-dorsally (Pita et al. 2015).

Measurements were taken during defined interaction periods. The beginning of an interaction period represented any approach made by the zebrafish towards the computer screen (Kalueff et al. 2013). In zebrafish, the turning angle represents a change in the directional heading (Kalueff et al. 2013), and we used it to define the end of each interaction period. Specifically, we measured the separation distance of the zebrafish prior to the frame where the zebrafish altered its directional heading beyond 45°, which in many cases, resulted in the zebrafish turning away from the computer screen. For each interaction period, we measured the separation distance (i.e., distance between the eye of the zebrafish and the animated conspecific) and interaction duration (i.e., difference between the start and end time of interaction period).

The zebrafish-specific cues that we were interested in were the eye and the horizontal striping pattern. The eyes have been shown to be an important cue involved in social recognition and the initiation of shoaling behavior (Landgraf et al. 2016), while the striping pattern has been shown to be used as a reference mark to assess changes in conspecific movement when individuals are interacting in groups (Guthrie and Muntz 1993; Bone and Moore 2008). We modified the magnitude of the social cues by manipulating the eye diameter and width of the conspecific body stripes, which changed as a result of the size of the conspecific itself, through a 2-fold increase (large conspecific = 8 cm) or 2-fold decrease (small conspecific = 2 cm) in size relative to a size-matched (intermediate conspecific = 4 cm) zebrafish. Because we kept the relative eye diameter and width of the lateral body stripes constant in each conspecific size, the net result was that the eyes and stripes were easier or more difficult to visually resolve in the large and small conspecifics, respectively.

We modified the level of visual risk by manipulating the vertical spatial position of the conspecific along the water column. Under stress, zebrafish position themselves at low vertical spatial positions (Egan et al. 2009; Kalueff et al. 2013). Thus, our virtual conspecific swam at either 7.6 cm (high risk level) or 12.7 cm (low risk level) from the bottom of the tank.

We modified the perceived depth of the conspecific by manipulating the presence/absence of the visual horizon. Treatments with the horizon absent (low perceived depth) had a white background with the virtual conspecific swimming. Treatments with the horizon present (high perceived depth) had a virtual substrate, the default terrain of the anyfish program. With the presence of this virtual substrate, we intended to have a distant horizon that the zebrafish could use as a spatial reference to enhance the perception of depth. When the virtual horizon was present, the conspecifics were 8 cm and 3 cm above the horizon in high and low risk treatments, respectively.

In our analyses, we included whether the zebrafish was using the center of acute vision (i.e., high acuity region of visual field) or retinal periphery (i.e., low acuity region of visual field) as a potential confounding factor because zebrafish may utilize high acuity regions of the visual field to increase their uptake of social information. We measured the position (angle in 3D space) of the conspecific relative to the eye of the zebrafish (Pita et al. 2015). The distance between the eye of the zebrafish and the center of the conspecific was calculated from 2D measurements of the top and side view camera recordings (supplementary material, Figure A1). To estimate where the center of acute vision projected in the visual space of our experimental arena, we calculated both the horizontal (angle relative to the horizontal plane of the zebrafish) and vertical (angle relative to the vertical plane of the zebrafish) positions of the eye of the zebrafish. We then evaluated the position of the conspecific relative to the projection of the zebrafish’s center of acute vision utilizing a 95% confidence interval range based on the retinal topographic map of the density of retinal ganglion cells (Pita et al. 2015). If the conspecific was within the 95% confidence interval range, we recorded that the fish was visualizing it with its center of acute vision for that frame. If the conspecific was outside the 95% confidence interval, we deemed it as being visualized by areas of the retina with low visual acuity at that time frame (e.g., retinal periphery). These estimates assumed that the zebrafish did not move its eyes considerably, as we were unable to accurately identify eye movement in the recorded videos. Nonetheless, some studies have shown that the vestibulo-ocular reflex allows fish to maintain a stabilized eye position when they are swimming (Easter and Johns 1974; Schairer and Bennett 1986).

### Statistical Analysis

We ran general linear mixed models using a repeated measures design in SAS (version 9.4). The code is available in the supplementary material. We tested the effects of the magnitude of the social cues, the level of visual risk, and the perceived depth on separation distance and interaction duration. Across trials, there were 10.66 ± 0.63 (mean ± SE) interaction behaviors analyzed per trial per individual. Each interaction behavior totaled approximately 23.9 ± 0.40 frames (mean ± SE). We also added to the models the region of the visual field that zebrafish used when interacting with conspecifics (i.e. center of acute vision vs. periphery) and the order in which the individual zebrafish were exposed to the treatments as independent factors. We used t-tests to assess differences between levels of a factor. In addition to the main effects, we also analyzed two-way interaction effects (i.e. magnitude of the social cues x level of visual risk, magnitude of the social cues x perceived depth, and the level of visual risk x perceived depth) on the separation distance and interaction duration. The original models included the three-way interaction (magnitude of the social cues x level of visual risk x perceived depth) but did not turn out to be significant; so, we removed it to increase the power of the tests.

## Results

### Separation Distance

The magnitude and perceived depth of the social cues and the level of visual risk all significantly affected zebrafish separation distance (Table 1a). Zebrafish increased their separation distance as the magnitude of social cues increased (i.e., larger conspecific with larger eyes and wider body stripes; Figure 1a). Zebrafish decreased their separation distance when the level of risk was higher (i.e., conspecific at bottom of the water column) relative to low risk levels (Figure 1b). Zebrafish increased their separation distance when the horizon was present compared to when it was absent (Figure 1c). Finally, we also found significant differences in the separation distance depending on the location of the conspecific in the zebrafish’s visual field. Specifically, zebrafish increased their separation distances when the conspecific was viewed with high acuity regions of the visual field (i.e., center of acute vision) relative to low acuity regions of the visual field (i.e., retinal periphery) (Figure 1d).

**Table 1.** Effects of the magnitude of the social cue, the visual social risk, the perceived depth of the social cue, and the visual field region used to view the virtual fish on: (a) separation distance, and (b) interaction duration. Significant effects are marked in bold.

|  | F | df | P |
| --- | --- | --- | --- |
| **Social Cue Magnitude** | **22.68** | **2,18** | **< 0.0001** |
| **Visual Risk** | **6.82** | **1,9** | **0.028** |
| **Perceived Depth** | **17.30** | **1,9** | **0.002** |
| **Visual Field Region** | **21.92** | **1,9** | **0.001** |
| **Social Cue Magnitude x Visual Risk** | **4.34** | **2,16** | **0.031** |
| Social Cue Magnitude x Perceived Depth | 1.33 | 2,16 | 0.293 |
| **Visual Risk x Perceived Depth** | **7.89** | **1,9** | **0.020** |
| **Social Cue Magnitude x Visual Field Region** | **4.59** | **2,13** | **0.031** |
| Perceived Depth x Visual Field Region | 4.10 | 1,6 | 0.089 |
| **Visual Risk x Visual Field Region** | **7.41** | **1,6** | **0.034** |

(b) Interaction Duration
|  | F | df | P |
| --- | --- | --- | --- |
| Social Cue Magnitude | 1.07 | 2,18 | 0.364 |
| Visual Risk | 0.56 | 1,9 | 0.475 |
| Perceived Depth | 1.21 | 1,9 | 0.300 |
| **Visual Field Region** | **5.69** | **1,9** | **0.041** |
| **Social Cue Magnitude x Visual Risk** | **4.23** | **2,16** | **0.034** |
| Social Cue Magnitude x Perceived Depth | 1.08 | 2,16 | 0.364 |
| Visual Risk x Perceived Depth | 0.06 | 1,9 | 0.816 |
| Social Cue Magnitude x Visual Field Region | 0.85 | 2,13 | 0.449 |
| Perceived Depth x Visual Field Region | 1.47 | 1,6 | 0.270 |
| Visual Risk x Visual Field Region | 0.46 | 1,6 | 0.524 |

**Figure 1.**
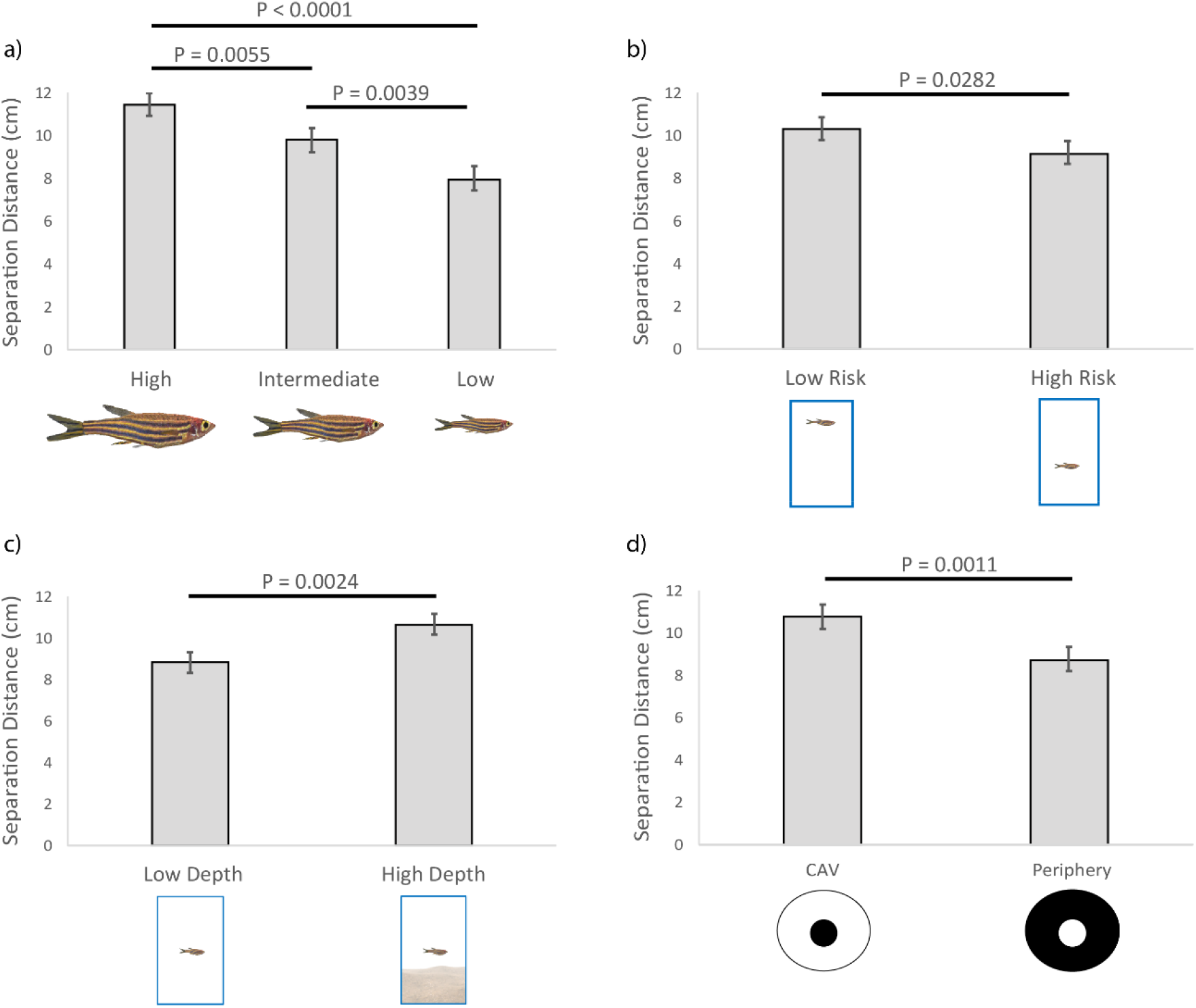
Variation in separation distance between the live and virtual fish (cm) relative to: (a) magnitude of the social cue (high, intermediate, low), (b) visual social risk (high risk, low risk), (c) perceived depth of the social cue (low, high), and (d) visual field region used to view the virtual fish (center of acute vision (CAV) and low acuity region (periphery)).

We also found two significant interaction effects between the factors of interest (Table 1a). First, the magnitude of social cues interacted with the level of visual social risk, whereby separation distances were significantly shorter in situations of high visual risk for conspecifics that had intermediate and low magnitude social cues (Figure 2a). However, no significant difference for separation distance was observed for visual risk with high social cue magnitude conspecifics (Figure 2a).

**Figure 2.**
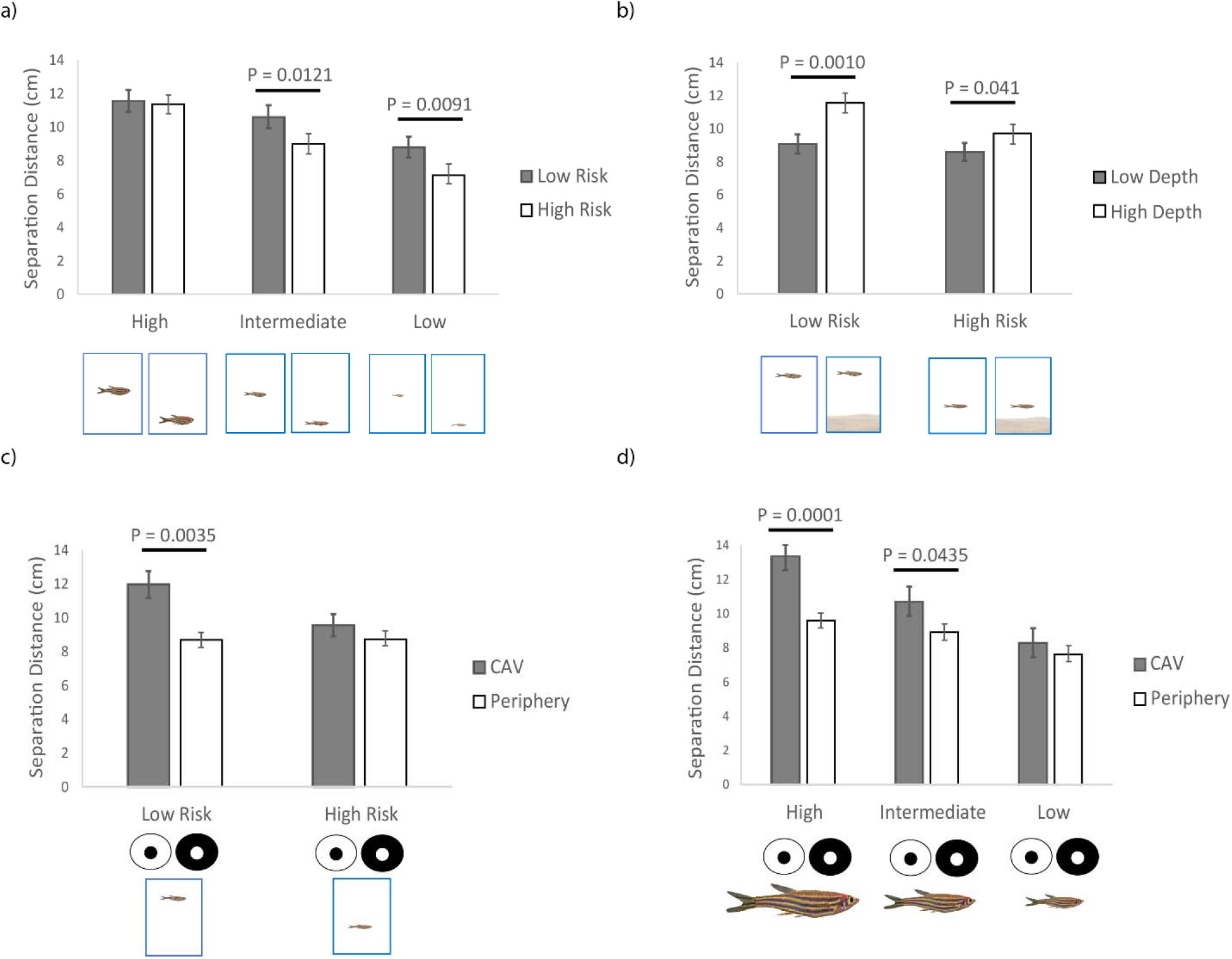
Variation in separation distance between the live and virtual fish (cm) relative to the simultaneous effect of: (a) magnitude of the social cue (high, intermediate, low) and visual social risk (high risk, low risk); (b) visual social risk and perceived depth of the social cue (low, high); (c) magnitude of the social cue and visual field region used to view the virtual fish (center of acute vision (CAV) and low acuity region (periphery)); and (d) visual social risk x visual field region used to view the virtual fish.

Second, the level of visual social risk also interacted with the perceived depth of the social cues such that in situations of high risk, the separation distance was longer when the horizon was present (high perceived depth) compared to when it was absent (low perceived depth). However, this difference in the separation distance became much more pronounced in situations of low risk (Figure 2b).

We also found two other significant interactions when considering the region of the visual field used during the interaction periods (Table 1a). First, the magnitude of the social cues interacted with the visual field region, such that separation distance was longer when the conspecific was viewed with high acuity regions of the visual field for conspecifics with large and normal magnitude social cues, however, no significant differences were detected for conspecifics with low magnitude social cues (Figure 2c). Second, the level of visual risk interacted with the visual field region, whereby separation distance was longer when the conspecific was viewed with high acuity regions of the visual field in low risk scenarios, but no significant difference was observed in high risk scenarios (Figure 2d).

### Interaction Duration

There were no significant effects of the magnitude and perceived depth of the social cues and the level of visual social risk, as single factors, on the duration of the interaction between the zebrafish and the conspecific (Table 1b). However, we found a significant interaction effect between the magnitude of the social cues and the level of visual social risk (Table 1b), by which interaction duration varied significantly across different social cue magnitudes (higher in intermediate and low relative to high magnitudes) only at high risk levels, but no significant difference in social cue magnitude at low risk levels (Figure 3a). Finally, we found a significant effect of the region of the visual field used during the interaction periods (Table 1b). Zebrafish spent more time interacting with conspecifics when conspecifics were viewed with the low acuity regions of the visual field compared to the high acuity regions (Figure 3b).

**Figure 3.**
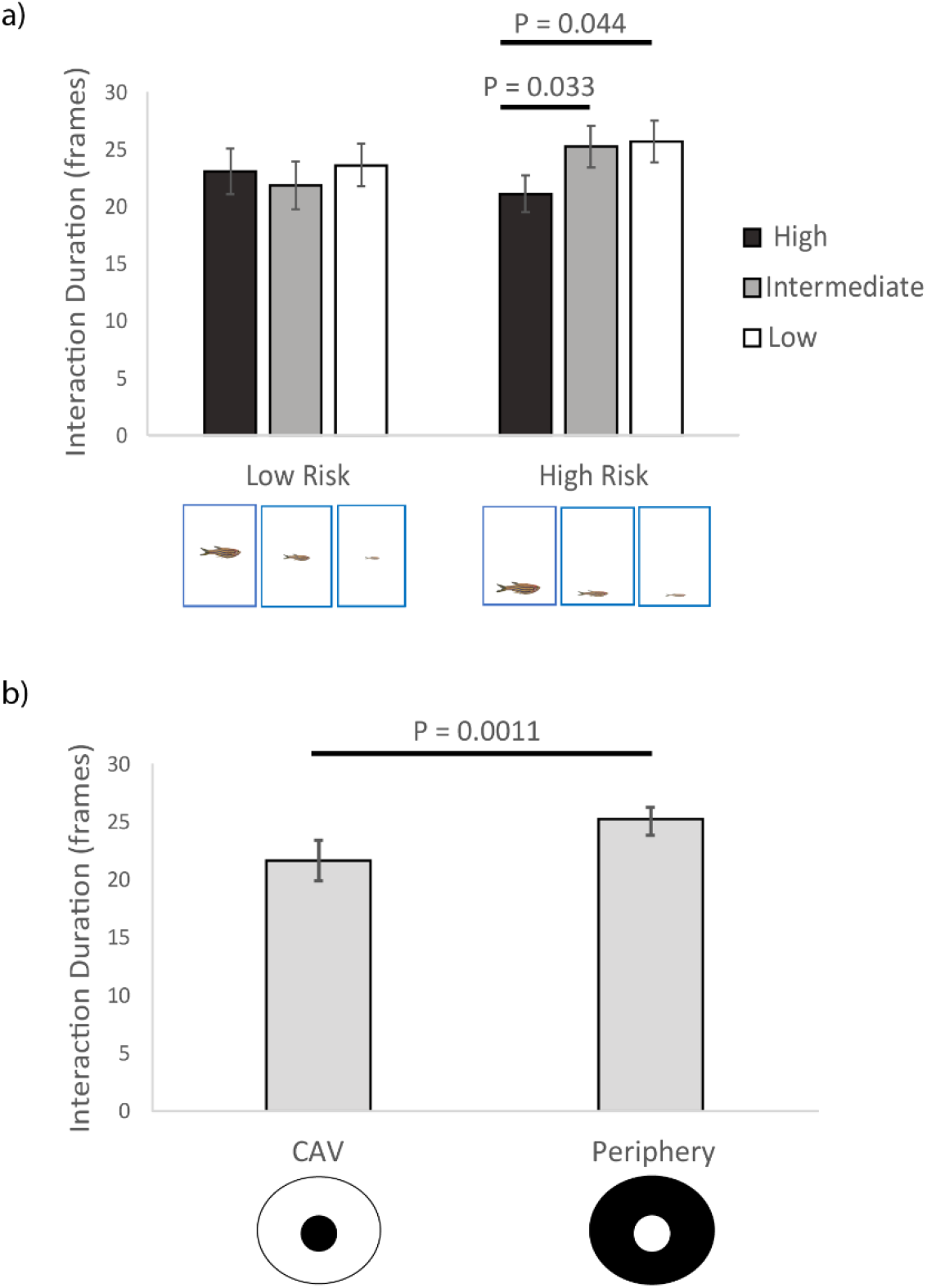
Duration (frames) of the interaction between the live and virtual fish relative to: (a) magnitude of the social cue (high, intermediate, low) and visual social risk (high risk, low risk); and (b) visual field region used to view the virtual fish (center of acute vision (CAV) and low acuity region (periphery)).

## Discussion

In a social pairwise scenario, zebrafish modified two aspects of their social behavior (spacing and time during a social interaction) as different features of the visual social environment changed (magnitude and perceived depth of the social cues, and level of visual social risk). Interestingly, zebrafish responded to our manipulations not just in an additive way but also in a multiplicative way. Zebrafish modulated their responses based on the combinations of features of their visual social environment such that their behavior towards a given feature depended on the levels of a second feature. These interaction effects were particularly pronounced for spacing behavior, but less so for interaction time.

All of our original predictions for single effects were supported by our findings. First, there was an inverse relationship between social cue magnitude and separation distance (i.e., zebrafish moved closer to the conspecific as the social cue magnitude decreased), which may allow for the improved resolution of the eye and body stripes that may be used to maintain social cohesion (Bone and Moore 2008). Second, zebrafish shortened their separation distances as the level of visual social risk increased likely because conspecifics at low vertical spatial positions were perceived as a higher risk cue (Kalueff et al. 2013). Third, when the virtual conspecific was displayed along with the horizon, zebrafish distanced themselves farther away probably because the referential information enhanced their ability to perceive depth and get a better estimate of neighbor distance (Layne et al. 1997; Caballero et al. 2015). However, the effects of these three features of the visual social environment cannot be interpreted in isolation because of the significant interaction effects we found.

The effects of visual social risk depended on the magnitude of the social cue. For intermediate and low social cue magnitudes, zebrafish decreased their separation distance during high risk compared to low risk situations. However, the effect of visual social risk vanished with high social cue magnitude. Considering that social cue magnitude scaled directly with the size of the conspecific, it is possible that these results could be explained by size disparity. In our experiment, live zebrafish were much smaller (i.e., ~3 cm difference in body length) than the high magnitude social cue conspecific, potentially leading to a decreased motivation to escalate a potential aggressive interaction had the live fish approached the conspecific (i.e., higher social costs). Other studies have shown that when given a choice, zebrafish prefer to associate size-matched or smaller conspecifics and avoid larger conspecifics (Fernandes et al. 2015; Bartolini et al. 2016). There is also the possibility that the high magnitude social cue conspecific could have been perceived as a predator; however, this is unlikely as we did not observe any antipredator associated behaviors (e.g., increased turning rate, jumping, erratic movement) (Kalueff et al. 2013).

The effects of the perceived depth of the social cue depended on the level of visual social risk. Specifically, we found greater neighbor separation distances when perceived depth was enhanced (horizon present) relative to when it was constrained (horizon absent) under situations of high social risk. However, this difference in separation distance relative to perceived depth was much more pronounced in the low social risk condition. It is possible that in the high risk situation, the effect of perceived depth is reduced as individuals shorten their separation distance in an attempt to increase the perception of social information or improve survival through dilution effects (Krause and Ruxton 2002).

In terms of interaction time, we did not find any significant single effect of any of the three features of the visual social environment we manipulated. Nevertheless, we did find a significant interaction effect between the magnitude of social cues and the level of visual social risk. Zebrafish spent less time interacting with high magnitude social cue conspecifics compared to intermediate and low magnitude social cue conspecifics in high social risk scenarios.

However, this social cue magnitude effect disappeared in low social risk scenarios. Considering that swimming speed is limited by body size (Aivaz and Ruckstuhl 2011), small individuals may be at a disadvantage when they interact with larger conspecifics in high risk environments due to their inability to maintain cohesion during evasive escape maneuvers (Peuhkuri 1998). The potential for small individuals to get left behind by larger group members may help explain their decreased motivation to interact with high magnitude social cue conspecifics under situations of high risk. Alternatively, individuals may also avoid grouping with larger conspecifics, especially when they maintain low vertical spatial positions, as in some fish species this has been associated with a higher likelihood to engage in aggressive behavior (Nakayama et al. 2009).

We also found that separation distance and interaction time varied depending on which portion of the visual field (high vs. low acuity) zebrafish used to view the conspecific. More specifically, zebrafish positioned themselves farther from, and interacted less when the conspecific was viewed with high vs. viewed with low acuity. This supports the idea that animals can resolve greater detail at farther distances when using retinal areas of high acuity, and can gather this high quality information faster during social interactions (Fernández-Juricic and Kowalski 2011). However, these single effects in the case of separation distance were modulated depending on the magnitude of the social cue and the level of visual social risk.

For example, the longer separation distance only occurred with large and normal social cue magnitudes when using high acuity relative to low acuity vision, but not with the small social cue magnitude. It is possible that when zebrafish interacted with smaller conspecifics, they may have changed their ability to choose between high vs. low acuity vision because of the visual challenges of resolving a small object. The ability of zebrafish to resolve a spatial grain is limited by the visual resolution of its eye. We used the spatial resolving power of zebrafish (Pita et al. 2015) to estimate whether or not individuals in our experiment would have been able to resolve the eyes and stripes of each of the social cue magnitudes based on their observed average neighbor distances (see supplementary material, Figure A3). We found that when interacting with intermediate and high social cue magnitude conspecifics (i.e., size-matched and larger conspecifics, respectively), individuals maintained a separation distance that would have allowed them to resolve key social cues (i.e., eye and stripes) with both high and low acuity vision (Figure A3). However, when interacting with low social cue magnitude conspecifics (smaller conspecifics), the separation distance they maintained would have allowed them to only resolve the eye and stripes with the high acuity vision but not with the low acuity vision. This suggests that the neighbor distances individuals maintained with small conspecifics may have limited their ability to resolve some key social cues, potentially constraining the types and intensities of their social interactions. Actually, evidence suggests that zebrafish take on a leader role when associating in groups with small conspecifics, maintaining positions at the front of the group where they can maximize their acquisition of personal (rather than social) information (Krause et al. 1998; Reebs 2001; Ward et al. 2002).

Finally, separation distance was longer when zebrafish used high relative to low acuity vision in the low visual social risk scenario, but the difference in the visual field region used vanished in the high social risk scenario. Birds have also been shown to vary their use of high vs. low acuity vision depending on perceived individual risk (i.e., exposure to a predator model vs. conspecific model; Butler and Fernández-Juricic 2018). In the case of zebrafish, under low perceived social risk, it is possible that individuals were able to divide their visual attention and adjust their spacing depending on which portion of their visual field they were using at the moment of visually inspecting the conspecific. However, these spatial adjustments relative to visual attention may be overridden by a situation of high social risk where the priority would be to minimize the chances of mortality (Pitcher and Parrish 1993; Krause and Ruxton 2002).

Overall, it appears that the visual social environment affects zebrafish social interactions in complex ways. Many vertebrates (fish and birds) interact socially in 3D space. Previous findings (Larsch & Baier 2018; Lemasson et al. 2018) and ours suggest that the rules underlying these 3D interactions are likely affected by multiple parameters acting simultaneously. In other words, the effects of some visual social parameters may depend on the intensity of other visual social parameters tuning up or down different social behavioral responses. We believe this is relevant because research on collective behavior sometimes tries to pin down simple rules governing between individual interactions (Schellinck and White 2011; Arganda et al. 2012; Pita et al. 2016). Although simplification is certainly necessary to understand complex behaviors, we suggest that considering the multiplicative effects of different visual components can enhance our ability to predict social interactions in groups.

One of the implications of the dynamic nature of zebrafish social behavior is that it could influence the spatial configuration of fish schools depending on habitat complexity and water turbidity. For example, under low perceived predation risk and with an horizon easily visible, individuals may maintain long separation distances, which would lead to low density groups with low intra-specific competition (Krause and Ruxton 2002). Additionally, low density fish schools could detect predators more quickly because of greater group visibility (Rountree and Sedberry 2009; Pita et al. 2015). In these low-density groups, individuals may orient themselves utilizing regions of the visual field with low acuity to monitor changes in conspecific behavior while the centers of acute vision may be focused on the environment for threats (Butler and Fernández-Juricic 2018). The presence of a threat could be quickly transmitted across the group via social information, leading to rapid changes in the spatial configuration of the school ending in spatially tighter (e.g., higher density) schools. More work along the lines of Lemasson et al. (2018) should establish the relative relevance of the different sources of visual social information in triggering these spatial structure changes as the number of group members increases (and their position in 3D space changes) in environments with different levels of habitat complexity and visibility.

## Acknowledgements

We are grateful to Deona Harris, Megan Bock, Elizabeth Brewer and Rachel Lim for helping in the video processing and data gathering.

## Compliance with Ethical Standards

The Institutional Animal Care and Use Committee of Purdue University (protocol 1207000675) approved all housing conditions and experimental procedures.

## Data Availability Statement

All data analyses during this study are published in the supplementary information of this article.

