## Supplementary Material for "The visual social environment affects non-additively neighbor spacing and interaction time in zebrafish"

***Experimental arena:***

We used two cameras to record the behavior of the live fish in response to the video playback of the virtual fish (Fig. A1).

**Figure A1**. Experimental arena and camera setup during video playback.

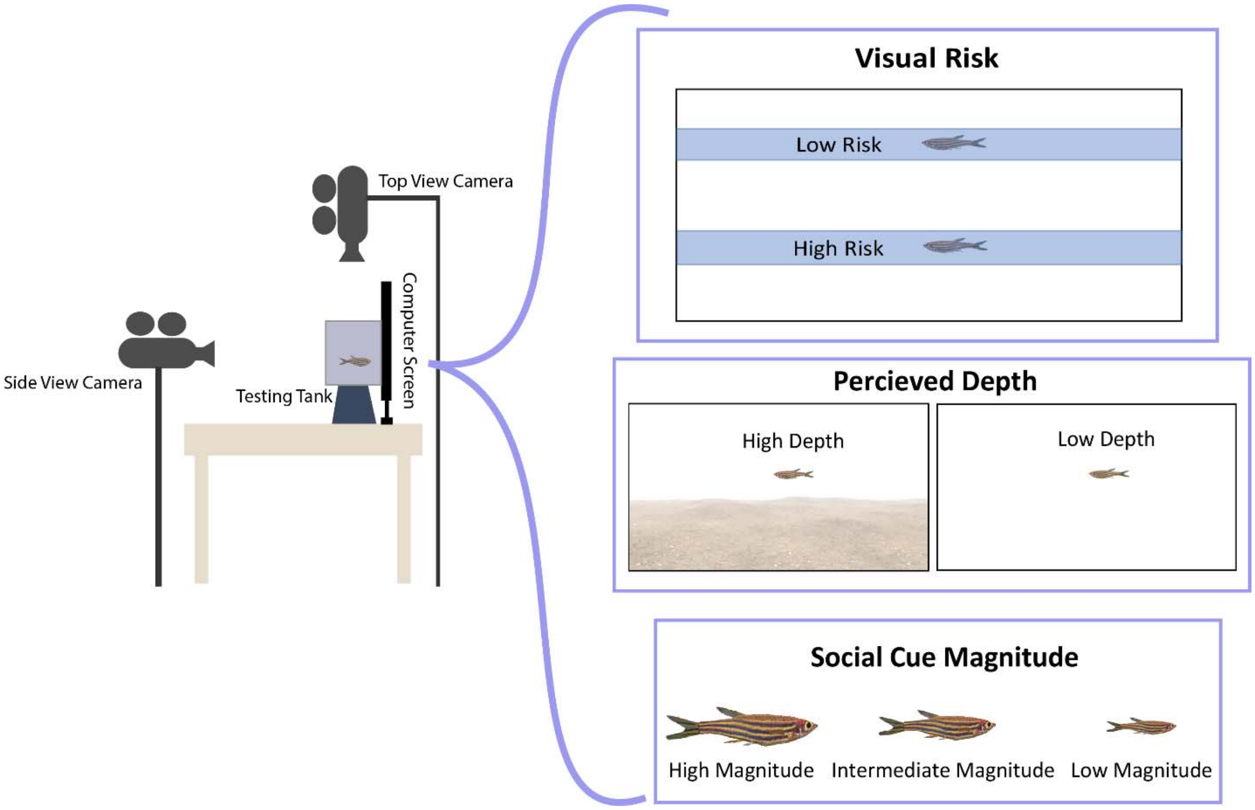


***Movement projections using automated tracking software***

**Figure A2**. Representative sample of the movement trajectory of the zebrafish (x and y position) before, during and after the virtual fish treatment exposure. Data for the figure was generated using the videotracking software idTracker (http://www.idtracker.es/) from the side view camera recording. Additional data available upon request.

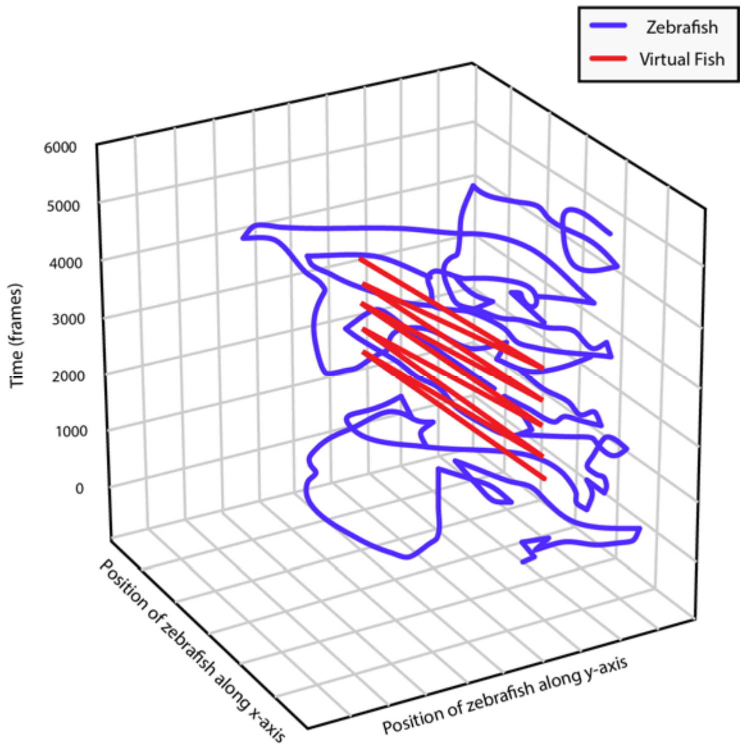


***Code used for the statistical analyses:***

We conducted our general linear mixed models using ProcMixed in SAS. The code used can be summarized as follows:

```
proc sort data=ddd; by trial id order;
proc mixed data=ddd;
class StimSize StimHeight Substrate CAVPER ID;
model DependentVariable* = StimSize  StimHeight Substrate CAVPER Order
StimSize*StimHeight  StimSize*Substrate StimHeight*Substrate StimSize*CAVPER
Substrate*CAVPER   StimHeight*CAVPER /solution outp=preds ddfm=bw;
repeated /subject=ID type=cs;
lsmeans StimSize  StimHeight Substrate CAVPER/diff;
proc plot data=preds; plot resid*pred;
proc univariate data=preds plot normal;  var resid;
run;
```

*Dependent variables were: Approach_Duration and Separation_Distance

Abbreviation of other factors:

- StimSize: the magnitude of the social cues (small, normal, large size of the virtual fish).
- StimHeight: the level of visual social risk (high, low position of the virtual fish in the water column).
- Substrate: the perceived depth of the social cues (presence, absence of the horizon).
- CAVPER: use of center of acute vision (high acuity vision) or retinal periphery (low acuity vision).
- ID: identity of the individual zebrafish.
- Order: order in which each individual zebrafish approached the virtual fish.

***Estimation of resolution limits:***

Visual acuity is positively associated with the density of retinal ganglion cells, which summate information from other retinal cell types into an electrical signal that is sent to the brain, and eye size (Pettigrew et al. 1988; Pettigrew and Manger 2008). Typically, higher visual acuity or spatial resolving power leads to a greater capacity to resolve objects of a finer scale at a given distance. Additionally, higher spatial resolving power allows animals to resolve objects from farther away. We estimated the distances at which zebrafish would be able to resolve the eye size and the stripe width of a conspecific (i.e., resolution limits) (Table 1). Retinal ganglion cell estimates of acuity were calculated from a previous paper (Pita et al. 2015), which were then incorporated into the following equation to calculate spatial resolving power: $\frac{RMF}{2}\sqrt{\frac{2D}{3}}$, where D represents the density of retinal ganglion cells and RMF represents the retinal magnification factor. The RMF was calculated as: $\frac{2\pi\ x\ PND}{360}$ , where PND is the posterior nodal distance, which is calculated by multiplying the radius of the zebrafish lens by 2.55 (Williams and Coletta 1987; Collin and Pettigrew 1988). Final spatial resolving power values are in units of cycles per degree. We estimated visual acuity using the retinal ganglion cell density of the center of acute vision (high acuity vision) as 1.89 cycles/degree, and the periphery of the retina (low acuity vision) as 0.81 cycles/degree. Actually, our low acuity vision estimates of spatial resolving power were similar to previous estimates based on photoreceptor densities (0.87 cycles/degree; Haug et al. 2010) and behavioral measurements (0.60 cycles/degree; Tappeiner et al. 2012; Cameron et al. 2013). Therefore, for the calculations in the next section we only used the ones we estimated from retinal ganglion cells (high and low acuity vision).

We then utilized the equation from (Tyrrell et al. 2013), $d = \frac{r}{\tan\frac{\alpha}{2}}$ , to calculate the maximum distance (d) that zebrafish could resolve conspecific social cues using the radius (r) of the eyes and stripes, with α being the inverse of the spatial resolving power. This equation assumes maximum visual contrast and optimal ambient light conditions.

We estimated the distances zebrafish would be able to resolve the eye and stripes of conspecifics with different sizes of the virtual fish and compared those distances to the averaged neighbor distances they maintained during the experiment (Figure A3). We considered resolution distances when the live fished used high acuity vision (Figure A3a) and low acuity vision (Figure A3b). These results suggest that zebrafish appeared maintain a distance that allowed them to resolve the eyes and stripes with high acuity vision for all virtual fish sizes (Figure A3a). However, the separation distance maintained by zebrafish would only allow for social cue resolution with low acuity vision for the large and normal virtual fish sizes, but not necessarily for the small virtual fish sizes (Figure A3b). Actually, when live fished interacted with the small virtual fish, they maintained a separation distance that would allow for the resolution of the eye and it was at the margin of the limits for resolving the stripes.

**Figure A3.** Distance at which zebrafish could resolve the eye and stripes of a conspecific (gray bars) relative to the averaged actual separation distance measured during the experiment for three sizes of the virtual fish (large, normal, small) considering resolution limits calculated using (a) high acuity vision and (b) low acuity vision.

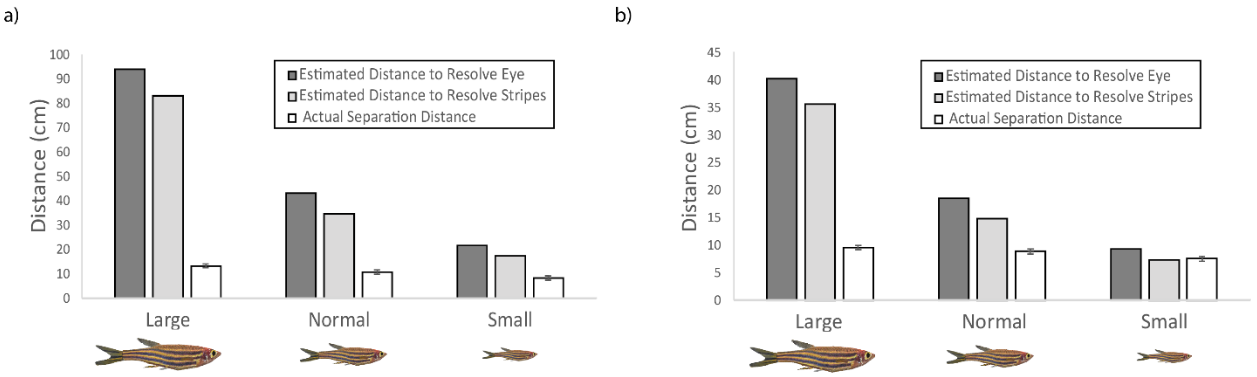
